# Improving science communication and organization visibility through Wikipedia: A case study of the American Association for Anatomy

**DOI:** 10.1101/2025.04.15.648547

**Authors:** Michael A. Pascoe

## Abstract

Wikipedia is one of the most widely accessed sources of scientific information globally, serving as a critical platform for public understanding and engagement with science. Despite its influence, many scientific societies are underrepresented or insufficiently described on the platform, limiting their visibility, outreach potential, and opportunities for public scholarship. This gap is particularly significant given Wikipedia’s high placement in search engine results and its role in shaping perceptions of scientific institutions. To address this issue, a structured case study was conducted focusing on the Wikipedia article for the American Association for Anatomy (AAA). A baseline evaluation revealed major deficiencies in article structure, historical coverage, citation quality, and alignment with Wikipedia’s editorial standards. Using a methodical editing process informed by science communication principles and content guidelines, the article was substantially revised. High-quality secondary sources were used to develop new sections on AAA’s mission, governance, publications, awards, meetings, and outreach efforts. Following these edits, the article’s classification improved from Stub to Start-class, its visibility and connectivity within Wikipedia increased, and the number of internal and external references, links, and structural elements grew substantially. This outcome demonstrates that scholarly engagement with Wikipedia can meaningfully enhance institutional presence and public accessibility of scientific knowledge. This case study offers a transparent and replicable model for scientific organizations and educators seeking to improve digital visibility and contribute to open-access science. It also highlights the potential for Wikipedia editing to serve as a form of public scholarship and a high-impact educational practice within the anatomical sciences.

## INTRODUCTION

### Science Communication Gap

There is currently a significant gap between scientific knowledge and public understanding, commonly referred to as the science communication gap.^1^ Much of the knowledge produced by the scientific community largely remains inaccessible to the general public due to paywalls, technical language, and limited reach of academic journals.^2,3,4^ As a result, important scientific insights frequently fail to reach broader audiences who could benefit from or contribute to their understanding.^5^

### The Free Encyclopedia That Anyone Can Edit

Wikipedia, launched in 2001 by Jimmy Wales and Larry Sanger, has become one of the most accessible and widely used sources of information in the world.^6^ Consistently ranking at the top of search engine results^7^, it receives billions of visits each month^8^, making it a critical entry point for the public seeking information on virtually any topic. Studies have shown that Wikipedia provides up-to-date and accurate summaries of scientific subjects, with accuracy levels comparable to those of traditional encyclopedias.^9^ Its influence has only grown with the rise of artificial intelligence and large language models, many of which rely on Wikipedia as a foundational source of training data.^10^ Despite growing recognition of Wikipedia’s value in science communication, little is known about how scientific organizations themselves are represented on the platform. Particularly in terms of the completeness, accuracy, tone, and accessibility of their Wikipedia pages.

### The American Association for Anatomy and the Role of Wikipedia

The American Association for Anatomy (AAA), founded in 1888, is dedicated to advancing anatomical science through research, education, and professional development.^11^ Scientific associations like the AAA play a critical role in fostering public engagement, not only by supporting their members but also by facilitating science communication and building trust with the public through outreach programs and accessible resources.^12^ For a scientific organization like the AAA, maintaining a high-quality Wikipedia page is crucial for several reasons. First, a well-crafted article enhances the organization’s credibility, serving as a trusted, verifiable source of information about its mission, history, and contributions to the field. This positions the AAA as an authoritative voice within the scientific community. Additionally, because Wikipedia is among the most visited websites globally, a strong presence on the platform significantly increases the visibility of the AAA’s initiatives, research, and educational programs to a diverse audience, including students, researchers, policymakers, and potential collaborators.^13^ Beyond visibility, the page also functions as a valuable tool for public outreach, helping communicate the importance of anatomical science and the AAA’s role within it to both specialists and the general public. A well-maintained Wikipedia page can also facilitate networking and collaboration by clearly presenting the organization’s activities and impact, potentially attracting new partnerships and funding opportunities. Moreover, it serves as a useful educational resource for educators, students, and professionals interested in anatomical sciences. Finally, due to Wikipedia’s search engine prominence^14^, a comprehensive and well-structured article can improve the AAA’s online discoverability, directing more users to its official resources. Despite these potential benefits, it remains unclear whether the current version of the AAA’s Wikipedia page is sufficiently developed to fulfill these important functions.

To bring clarity to this issue, a systematic evaluation of the current Wikipedia article on the AAA was conducted. Using criteria grounded in best practices for science communication, public engagement, and Wikipedia’s own content guidelines, the page’s accuracy, completeness, tone, and accessibility were assessed. The findings reveal key deficiencies, particularly in historical coverage, clarity of mission, and representation of the AAA’s ongoing activities, that likely limit the page’s effectiveness as a public-facing resource. These results suggest that improving organizational Wikipedia entries is a practical and impactful way to enhance science communication, public trust, and institutional visibility. A secondary purpose of this project was to document the process of updating the AAA’s Wikipedia entry in a transparent and reproducible manner, offering a potential model for other scientific organizations seeking to improve their public presence through Wikipedia.

## DESCRIPTION

### Baseline Evaluation

Prior to initiating any edits, the existing Wikipedia article titled American Association for Anatomy was systematically reviewed to establish a baseline for comparison and improvement. The version assessed was last edited on November 29, 2024^15^ and was accessed on April 4, 2025 (Figure 1). This version of the article was critically evaluated against several criteria drawn from the Wikipedia Manual of Style.^16^ These criteria included article structure and organization, citation quality and quantity, gaps in content or outdated information, and the presence of bias, promotional tone, or unverified claims.

**Figure 1.**
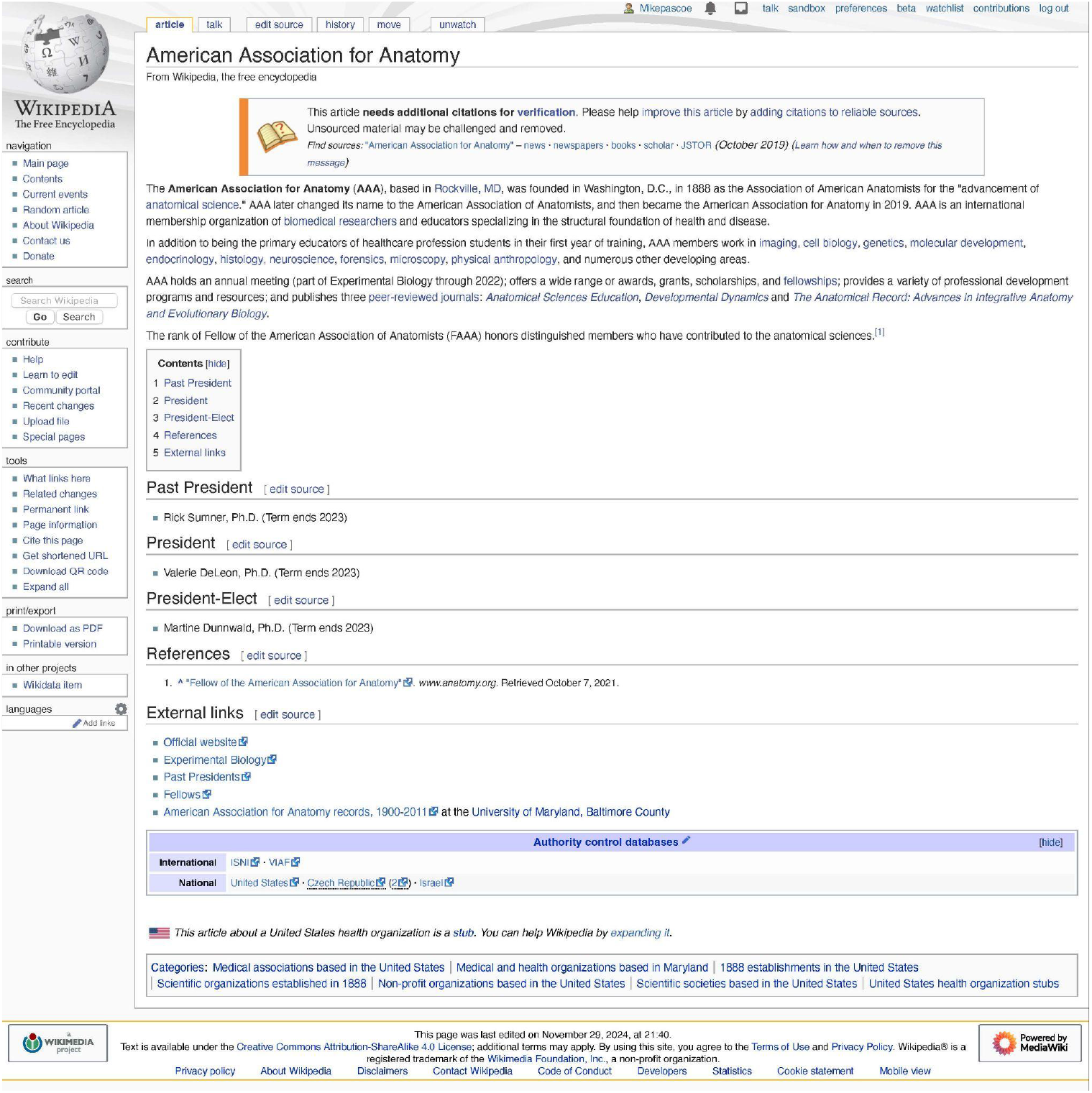
Screenshot of the Wikipedia page for the American Association for Anatomy (AAA) taken on the baseline analysis date. Note the cleanup tag “More citations needed” at the top of the article.

To assess the state of the Wikipedia article for the AAA, a structured set of evaluation metrics was employed across five domains: content quality, engagement and visibility, edit history and community involvement, technical and structural completeness, and word count analysis. These metrics reflected the article’s current state and its improvement potential.

The content quality of the article was limited at baseline. It was rated as a “Stub” by associated WikiProjects, indicating that the article contained minimal content and lacked comprehensive coverage of the topic. According to Wikipedia’s content assessment scale^17^, a Stub-class article offers only a minimal description of a topic and fails to meet the criteria for Start-class designation. Such articles typically lack meaningful content, may resemble dictionary definitions, and provide little context to help readers understand the topic’s significance. For readers, this often results in a limited and underdeveloped experience. Improving a Stub-class article involves adding substantive, well-referenced information, particularly content that highlights the topic’s relevance or importance, thereby helping it progress toward Start-class quality. The Talk Page showed no active discussions, suggesting that the article had not yet drawn the attention of the editing community for collaborative development. Only one WikiProject (WikiProject United States) had formally adopted the article, pointing to limited topical or disciplinary engagement. The article included a single internal reference and no external citations, meaning 100% of its sourcing was derived from within Wikipedia itself. This absence of external references undermines the article’s verifiability and adherence to Wikipedia’s reliable sourcing guidelines. The article featured 18 internal links to other Wikipedia pages, which supports basic navigational integration. At the time of review, the article was flagged with the cleanup tag “More citations needed” indicating that substantial portions lacked sufficient references to reliable sources. This warning suggests the potential for unsourced material to be challenged or removed, underscoring the need for improved citation practices. The article also lacked images or tables entirely, limiting its visual appeal and utility as an informative resource.

In terms of engagement and visibility, the article had been viewed a total of 4,067 times, with a daily average of approximately one pageview (Pageviews Analysis Tool^18^; Figure 2). This relatively low level of traffic might reflect both the article’s short length and its limited discoverability. The “What links here” tool revealed the article had 28 direct inbound links (i.e., links from other Wikipedia pages that reference it explicitly) and 53 indirect inbound links, indicating a modest degree of internal connectivity. See Appendix for a comprehensive list of inbound links. Beyond Wikipedia, five unique external websites linked to the article. In Google search results, the article ranked fifth when searching for the organization’s name, a respectable position that nonetheless suggests there is room to improve search engine optimization through enhanced content and metadata.

**Figure 2.**
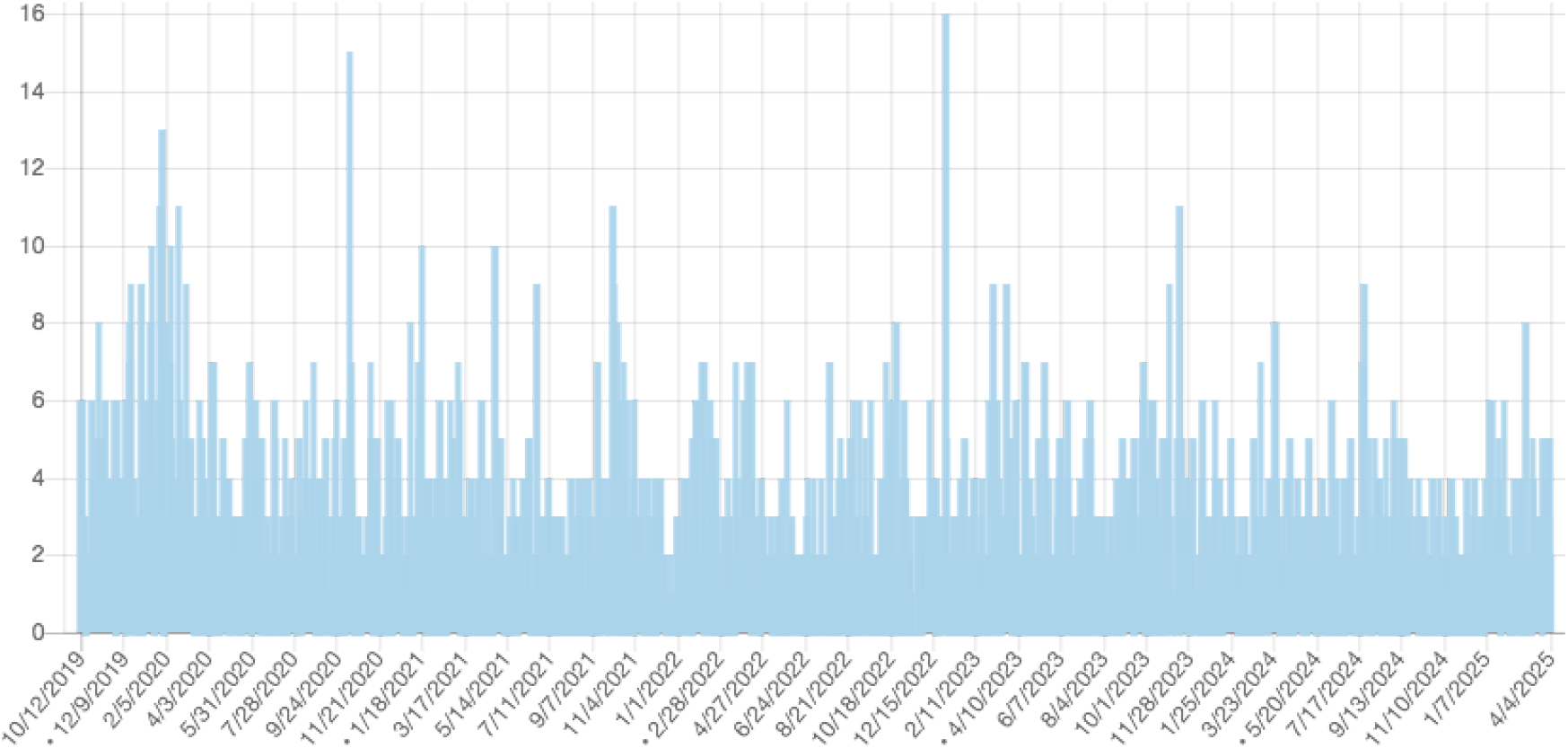
Daily Wikipedia pageview counts for the American Association for Anatomy from October 2019 to April 2025, illustrating trends in public interest leading up to the baseline analysis date (April 4, 2025).

The article’s edit history showed 55 total revisions made by 17 unique editors, indicating some degree of interest over time but no sustained editorial activity. The article’s edit history showed that it was originally created on February 12, 2009, and had seen limited activity since. No substantive updates had been made in recent years, with the most recent edit made on November 29, 2024. The article size was 3,003 bytes, consistent with its Stub-class designation. Fewer than 30 users were watching the page, which is typical for low-traffic articles but reflects limited ongoing editorial surveillance or interest from the Wikipedia community.

From a technical and structural standpoint, the article lacked a standardized infobox, which would normally summarize key facts and improve both visual structure and searchability. It was, however, linked to a Wikidata entry (Q4743046)^19^, establishing a connection to structured data within the larger Wikimedia ecosystem. The article was assigned to seven categories and consisted of five main sections, reflecting some effort at organization but not enough to provide depth or breadth in coverage.

A word count analysis via a publicly available, user-developed script^20^ further confirmed the article’s brevity. The main body contained only 219 words, and the references section contained 74 words—largely metadata or citations to internal Wikipedia pages. This word count is far below the level typical of comprehensive articles on notable academic or professional organizations.

Together, these baseline metrics reveal an article that, while established, remained underdeveloped across nearly all dimensions. They served as a reference point for efforts undertaken in this project to improve the article’s quality, visibility, and utility as a digital resource for both general readers and those within the anatomical sciences.

To aid in determining appropriate content and structural improvements, a comparable professional society’s Wikipedia article, The American Physiological Society^21^, was identified and assessed across the same five domains.

### Editing and Improvement

Following the baseline analysis, an editing strategy was developed to address the structural, informational, and citation-related gaps identified in the existing American Association for Anatomy Wikipedia article. This strategy was informed by the format and content of a comparable professional society entry, the American Physiological Society. Key elements of the improvement plan included expanding the article’s outline, sourcing high-quality secondary references, and drafting new content within the author’s personal Wikipedia user sandbox to ensure compliance with Wikipedia’s editorial standards before public publication.

The existing article text was first copied into the sandbox, where edits were made iteratively over a ten-day period (April 5–15, 2025; Table 1). Proposed additions were based on a curated list of reliable sources, including the official AAA website, peer-reviewed publications, historic newspaper archives, university records (e.g., University of Maryland, Baltimore County), books, news articles, press releases, and content from databases like GuideStar. Emphasis was placed on using high-quality secondary sources to maintain verifiability and align with Wikipedia’s core policies: a neutral point of view, no original research, and adherence to verifiability. Tables with information about presidents and locations of annual meetings were added to the article to improve readability, highlight key facts, and meet Wikipedia’s formatting standards.

**Table 1.** Detailed edit history of the Wikipedia article on the American Association for Anatomy (AAA) from April 5 to April 11, 2025. The table documents individual contributions, including timestamp, editor username, changes in page size, and the nature of content additions or modifications. The data illustrates the collaborative process and progressive development of the article leading up to its enhanced version.

| Edit Time | Edit Date | Contributor Username | Page Size (Bytes) | Bytes Changed | Edit Summary Provided by User |
| --- | --- | --- | --- | --- | --- |
| 8:12 | 5-April-2025 | Mikepascoe | 3051 | 3005 | Copied existing article text |
| 6:11 | 6-April-2025 | Mikepascoe | 7963 | 4912 | Added short description, organization infobox, categories |
| 7:11 | 6-April-2025 | Mikepascoe | 8035 | 72 | Added info to infobox |
| 7:31 | 6-April-2025 | Mikepascoe | 8148 | 113 | Added more info to infobox |
| 11:14 | 6-April-2025 | JJMC89 bot | 8084 | -64 | Removed WP:NFCC violation(s). Non-free files are only permitted in articles |
| 8:50 | 7-April-2025 | Mikepascoe | 10,293 | 2209 | Added more info to infobox and rewrite of introduction / lead section |
| 9:22 | 7-April-2025 | Mikepascoe | 13,864 | 3571 | History section written |
| 9:26 | 7-April-2025 | Mikepascoe | 14,424 | 560 | Mission Vision Values section written |
| 9:35 | 7-April-2025 | Mikepascoe | 14,452 | 28 | Added revenue and expenses to infobox (sourced from GuideStar) |
| 9:44 | 7-April-2025 | Mikepascoe | 16,636 | 2184 | Governance and Structure: Section written |
| 9:58 | 7-April-2025 | Mikepascoe | 20,528 | 3892 | Publications section written |
| 10:05 | 7-April-2025 | Mikepascoe | 22,449 | 1921 | Annual meeting section written |
| 10:13 | 7-April-2025 | Mikepascoe | 24,919 | 2470 | Awards and Recognition section written |
| 10:14 | 7-April-2025 | Mikepascoe | 24,915 | -4 | History: corrected original association name |
| 10:50 | 7-April-2025 | Mikepascoe | 27,193 | 2278 | Educational and Outreach Initiatives section written |
| 10:55 | 7-April-2025 | Mikepascoe | 29,141 | 1948 | Membership section written |
| 10:58 | 7-April-2025 | Mikepascoe | 29,397 | 256 | See Also section written |
| 11:02 | 7-April-2025 | Mikepascoe | 30,177 | 780 | External Links added, EB removed as there is no longer an active affiliation |
| 13:10 | 7-April-2025 | Mikepascoe | 30,317 | 140 | History: added Photo of Joseph Leidy |
| 13:14 | 7-April-2025 | Mikepascoe | 30,535 | 218 | Added more categories |
| 13:15 | 7-April-2025 | Mikepascoe | 30,430 | -105 | Removed two Categories that were redlinks |
| 13:36 | 7-April-2025 | Mikepascoe | 31,515 | 1085 | Added info from the 2024 AAA Year in Review report PDF |
| 13:46 | 7-April-2025 | Mikepascoe | 33,715 | 2200 | Added several news stories |
| 13:47 | 7-April-2025 | Mikepascoe | 33,876 | 161 | Added photo of Florence Sabin |
| 13:51 | 7-April-2025 | Mikepascoe | 34,320 | 444 | Added reference to John E. Pauly book |
| 8:25 | 8-April-2025 | Mikepascoe | 34,712 | 392 | Membership: Added membership distribution data from 2024 year in review report |
| 8:43 | 8-April-2025 | Mikepascoe | 34,586 | -126 | Cleaned up errors in References |
| 15:38 | 8-April-2025 | Mikepascoe | 37,527 | 2941 | Added more examples of public engagement activities from the Dunnwald et al 2025 article |
| 15:51 | 8-April-2025 | Mikepascoe | 38,334 | 807 | History: added information found on the UMBC webpage |
| 9:17 | 9-April-2025 | Mikepascoe | 38,524 | 190 | Added scan of 1888 newspaper announcement of founding meeting of the AAA |
| 9:39 | 9-April-2025 | Mikepascoe | 43,239 | 4715 | Past presidents table added, wikilinks added to names of presidents |
| 9:45 | 10-April-2025 | Mikepascoe | 43,360 | 121 | Added caption to presidents table |
| 10:11 | 10-April-2025 | Mikepascoe | 44,614 | 1254 | Added annual meetings table |
| 11:10 | 10-April-2025 | Bearcat | 44,113 | -501 | Categories removed, cannot be in draft space |
| 13:36 | 10-April-2025 | Mikepascoe | 44,552 | 439 | Social media links added to Educational and Outreach Initiatives section |
| 13:56 | 10-April-2025 | Mikepascoe | 45,489 | 937 | Awards and Recognition: Added details about the AJ Ladman award |
| 13:58 | 10-April-2025 | Mikepascoe | 45,783 | 294 | History: Added info about the support of the AACA |
| 14:00 | 10-April-2025 | Mikepascoe | 46,121 | 338 | Annual and Regional Meetings: added details about member engagement from Keith Moore interview |
| 13:27 | 11-April-2025 | Mikepascoe | 47,381 | 1260 | Black in Anatomy partnership added to Educational and Outreach Initiatives section |
| 13:35 | 11-April-2025 | Mikepascoe | 47,589 | 208 | Updated President positions and clarified composition of Board of directors in the infobox |
| 20:28 | 12-April-2025 | Mikepascoe | 47,754 | 165 | Added scan of page from 1906 issue of The Anatomical Record |
| 10:12 | 15-April-2025 | Mikepascoe | 48,218 | 27 | Added logo to infobox using appropriate Non-free use rationale |
| 10:32 | 15-April-2025 | Mikepascoe | 48,274 | 56 | Educational and Outreach Initiatives: Added flickr social media platform link |

Substantial content was added to broaden the article’s coverage. Entirely new sections were developed, including: History, Mission, Vision, and Values, Governance and Structure, Publications, Annual and Regional Meetings, Awards and Recognition, Educational and Outreach Initiatives, and Membership. The References and External Links sections were also significantly expanded to support these additions. Where appropriate, photographs, images, and logos (e.g., the AAA logo) were incorporated using Wikimedia Commons and fair use protocols. Additionally, an advanced search was conducted to identify other articles referencing the “American Association for Anatomy.” Where appropriate, internal links (i.e., Wikilinks) to the updated AAA article were added to enhance visibility and connectivity across the encyclopedia. As a result of these changes, the article expanded from 219 to 3,109 words in the main body, with reference word count increasing from 74 to 813, and the number of internal links rising from 18 to 107.

To ensure transparency and collaboration with subject matter stakeholders, the AAA communications team was contacted. The purpose of the editing project was explained, and an invitation to contribute was extended. AAA responded positively and agreed to assist by providing images. A list of suggested photographs was shared along with guidance on how to release them in accordance with Wikimedia licensing standards. In addition, to broaden awareness and encourage further collaboration, a message was posted on April 15, 2025, to AAA member colleagues via the official Anatomy Network discussion platform. The post summarized the goals of the Wikipedia editing initiative and invited members who speak languages other than English to help translate the updated American Association for Anatomy article. This outreach aimed to support the global dissemination of anatomy-related knowledge and promote multilingual access to accurate information about the Association.

Prior to publishing the updated content, a notification summarizing proposed changes was posted on the article’s talk page on April 8, 2025. This message invited feedback from other Wikipedia contributors and provided an opportunity for community input. Additional WikiProject banners (Anatomy, Medicine, and Organizations) were added to the talk page to encourage collaboration and signal the article’s relevance to broader interest groups within Wikipedia.

After a one-week waiting period, final edits were made in response to user feedback, and the revised content was transferred from the sandbox to the main article on April 15, 2025.

Finally, the article’s category tags were reviewed and updated to reflect its enhanced scope. As of April 4, 2025, the article included seven categories, such as Medical associations based in the United States and Scientific societies based in the United States. A comparison with the categorization of similar societies (e.g., the APS; Wikipedia article created March 10, 2009) helped ensure consistency and accuracy in classification. Article characteristics were documented through a summary table (Table 2), facilitating comparison with baseline data and providing a framework for future analysis of impact. A screenshot of the edited article at the time of submission is provided in Figure 3. The data highlight the transformation of the article from a minimal Stub into a substantially enriched Start-class entry, closely approaching the level of peer organizations such as the American Physiological Society.

**Table 2.**
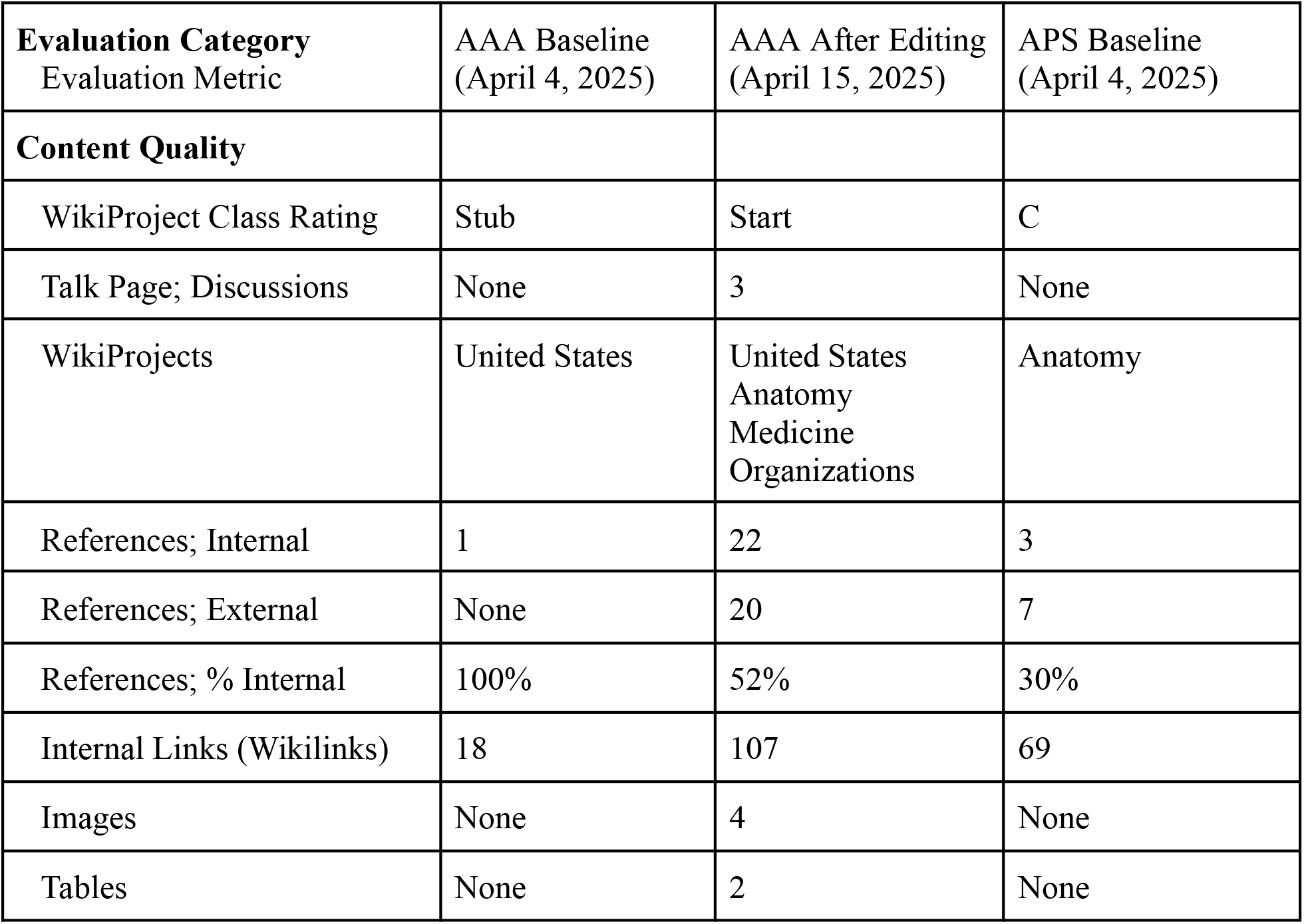
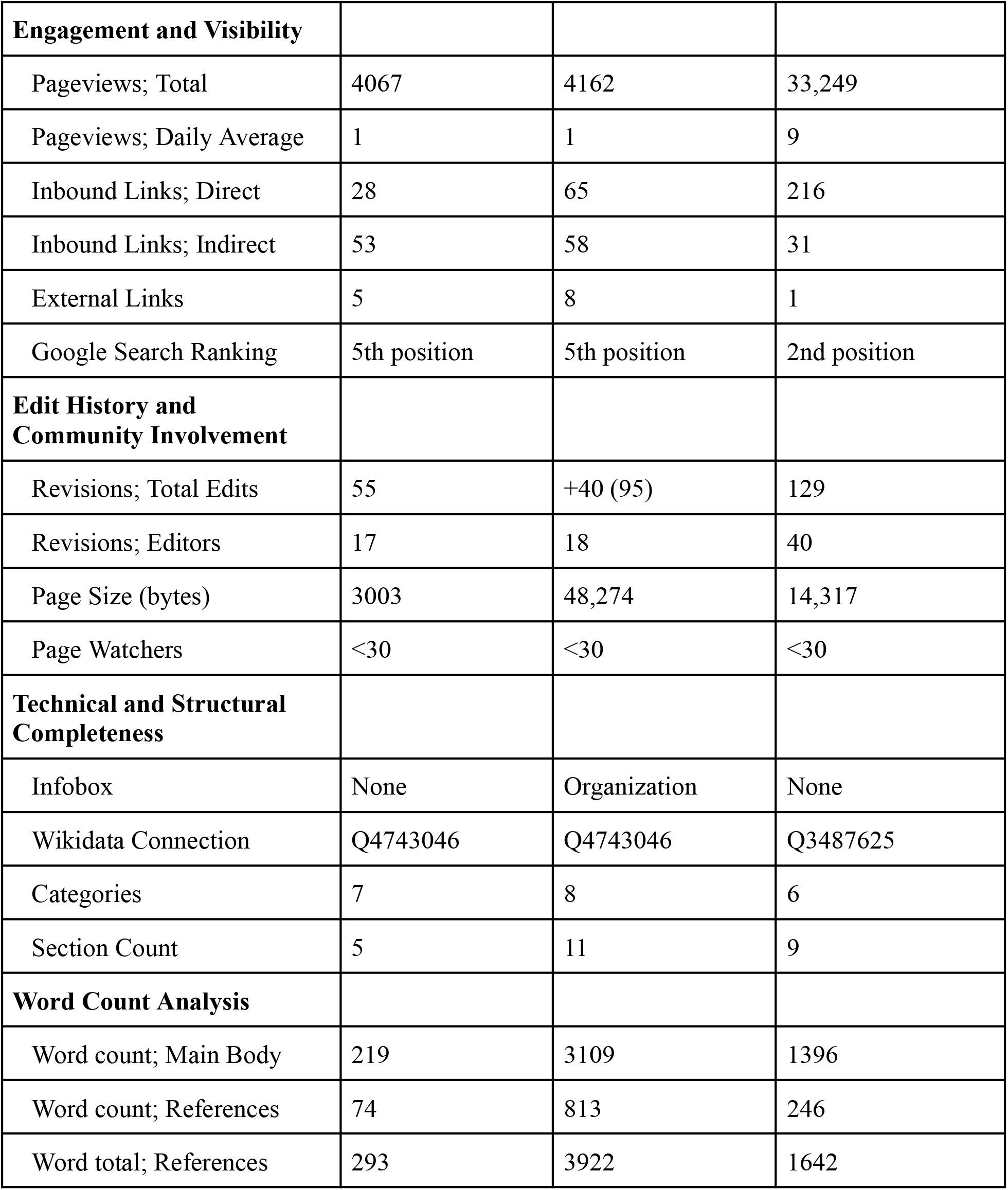
Comparison of Wikipedia article evaluation metrics for the American Association for Anatomy (AAA) before and after the editing initiative (baseline: April 4, 2025; post-editing: April 15, 2025), with American Physiological Society (APS) data included for reference.

**Figure 3.**
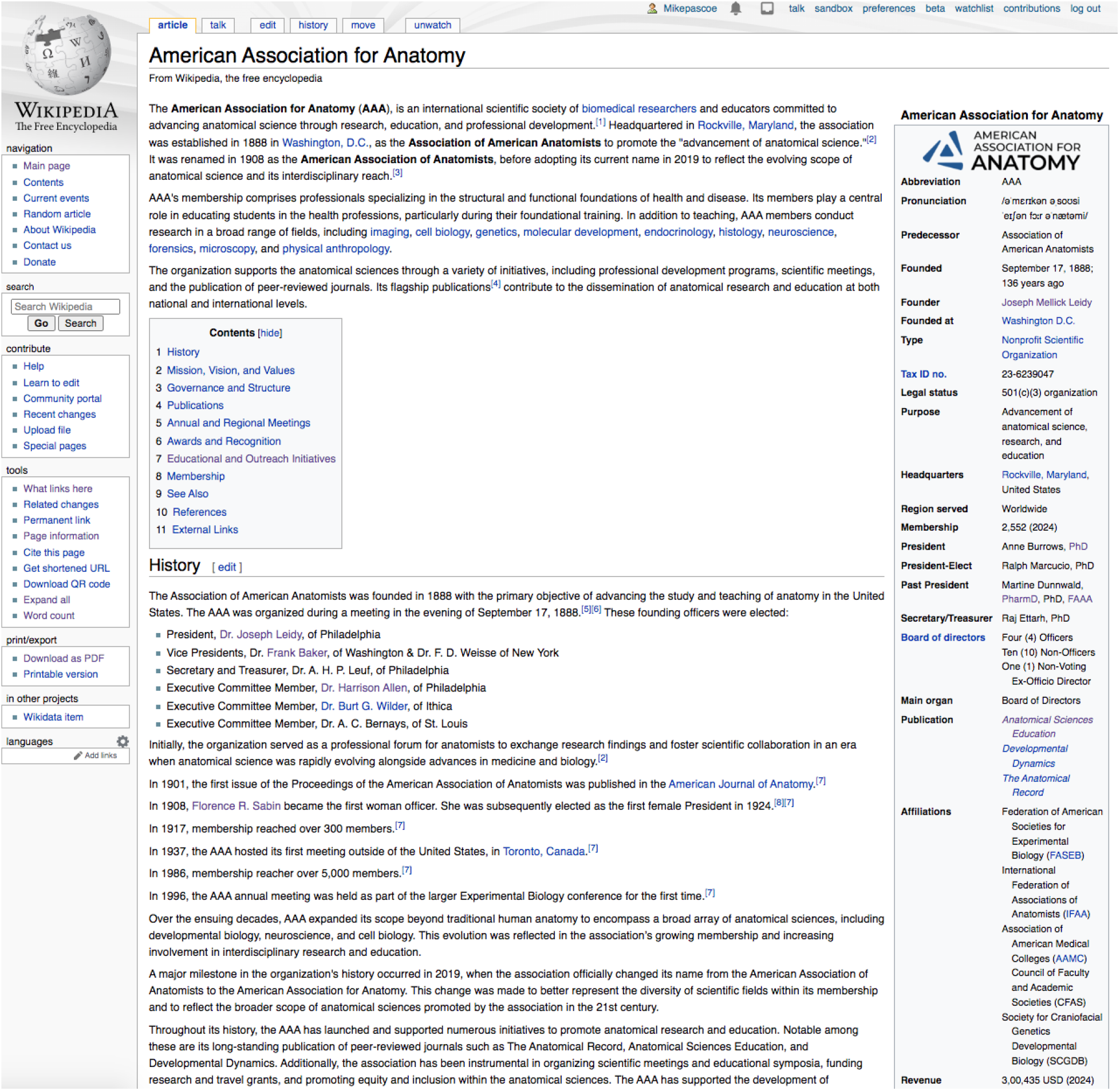
Screenshot of the Wikipedia page for the American Association for Anatomy (AAA) taken after the editing initiative (April 15, 2025). This screenshot captures only the top one-fifth of the entire article.

## DISCUSSION

This project underscores the educational and scholarly value of contributing to Wikipedia, particularly in the context of improving public-facing content about scientific organizations, such as the American Association for Anatomy (AAA). Wikipedia serves not only as a widely accessed reference platform but also as an active venue for public scholarship, where contributions by subject-matter experts can make a lasting impact on science communication and public understanding.^22^ By engaging with Wikipedia as a legitimate scholarly activity, academics can reach broader audiences while modeling best practices in information stewardship.

### Educational and Scholarly Value

Editing Wikipedia offers a valuable opportunity for scholarly engagement outside traditional publication models. It demands that contributors master skills aligned with academic values (e.g., such as rigorous sourcing, clear writing, and intellectual humility) while also navigating Wikipedia’s community norms. For educators, Wikipedia projects can serve as high-impact assignments that promote information literacy, collaborative authorship, and public engagement. For scholarly societies, these contributions enhance public visibility and reinforce their role as trusted authorities in their field. The improved AAA article now functions as both an outreach tool and a living, evolving record of the association’s activities and legacy. Its measurable improvement (e.g., increased references, added visual elements, enhanced article structure, and upgraded classification) demonstrates the value of strategic Wikipedia engagement.

### Barriers and Lessons Learned

Several barriers were encountered during the editing process. Chief among these was identifying high-quality, secondary sources that met Wikipedia’s standards for reliability and verifiability. Much of the relevant content about the AAA (e.g., historical information) is found in primary documents or internal communications, which are typically discouraged on Wikipedia. The need for neutrality also posed challenges, requiring careful editing to avoid promotional tone and to ensure balanced representation. Additional formatting limitations, such as restrictions on logo usage under fair use, also required creative problem solving, particularly in sourcing freely licensed images. The absence of existing high-resolution, appropriately licensed media highlighted the importance of proactive engagement with organizational communications teams to improve Wikimedia Commons contributions.

Although content disputes were limited in this case, they represent a known challenge for academic contributors. These may stem from differing interpretations of neutrality or the suitability of specific sources. Early posting of proposed edits on the talk page and adherence to Wikipedia’s content policies helped mitigate potential friction. Instructors and scholars embarking on similar efforts should anticipate these dynamics and approach them with collaborative intent.

### Future Directions

Looking forward, the AAA article could be nominated for the “Did You Know?” section of Wikipedia’s main page, an effective mechanism to increase article visibility and reach. A long-term aspiration is to develop the article to the point that it meets the strict criteria for nomination for Good Article status, and eventually, Featured Article status. These designations require rigorous adherence to Wikipedia’s standards and reflect well on the organization’s public profile.

To further enhance accessibility, future efforts could include translating the article into other languages, particularly those aligned with global anatomical society membership. Given the significant number of anatomy scholars and students in Asia, versions in Mandarin, Hindi, Korean, or Japanese could dramatically expand access. Such translation efforts would benefit from collaboration with multilingual anatomy educators and native speakers, combining subject expertise with cultural and linguistic fluency.

### Encouraging Participation in Anatomy Education

This project illustrates how Wikipedia engagement can be encouraged as part of formal anatomy education. Student and faculty contributions to anatomy-related articles promote scientific literacy, empower learners as knowledge producers, and help demystify the editorial process behind one of the world’s most-used information sources. Embedding Wikipedia assignments into coursework has already shown promise in health professions education^23^, and similar models could be applied within anatomy programs. Educators may draw on training resources such as those offered by Wiki Education to scaffold student participation.

Moreover, this project serves as an open invitation for others to participate in the continual improvement of the AAA article and related content, i.e., no contribution is too small. As Logan et al.^22^ argue, the public writing of science is not simply about dissemination, but about dialogue: small edits accumulate, forming a collective scholarly enterprise that reflects the community’s shared values and evolving knowledge base.

### Opportunities for Societies and Organizations

The AAA is not alone in having a limited Wikipedia presence. A review of Wikipedia articles related to international anatomical societies reveals significant opportunity for improvement. The Association of Clinical Anatomists (ACAA), International Federation of Associations of Anatomists (IFAA), British Association of Clinical Anatomists (BACA), and Anatomical Society (Anat Soc) have articles rated as Stub or have not yet been assessed. Others, such as the European Association of Clinical Anatomists (EACA) and The Australian and New Zealand Association of Clinical Anatomists (ANZACA) are completely absent.

There are currently 114 articles categorized under “Scientific societies based in the United States” on Wikipedia. Only 13% of these are rated as C-class, while the majority are Start-class (56%) or Stub-class (24%). This reveals a major opportunity for anatomy and other scientific societies to appraise the quality of their digital presence and to contribute systematically to Wikipedia improvement efforts.

Anatomy societies and institutions are well-positioned to advance public scholarship by actively supporting initiatives that enhance the quality and accessibility of anatomical knowledge^24^ on platforms like Wikipedia. They can host Wikipedia editing workshops at annual meetings to engage members directly in content improvement. Establishing structured internships or fellowships focused on public scholarship offers trainees meaningful opportunities to contribute while gaining valuable experience. Societies can also encourage their members to assess and enhance Wikipedia content related to their own organizations, promoting both accuracy and visibility. Additionally, developing guidance and media release protocols can facilitate the inclusion of high-quality images and data that are compatible with Wikipedia’s open-access standards. Partnering with groups such as Wiki Education or local Wikimedia affiliates can further amplify these efforts through training and outreach, helping to build a sustained culture of public engagement within the anatomical sciences.

### Conclusion

By bringing the American Association for Anatomy’s Wikipedia article up to a higher standard, this project demonstrates the impact and scholarly value of engaging with Wikipedia as a communication platform. It offers a replicable model for how academic contributors can apply their expertise in service of public knowledge, while also enriching their own understanding of open-access publishing, information ethics, and community authorship. There remains a rich opportunity for anatomy educators, students, and societies to join in the global effort to share knowledge freely, to reflect the evolving field of anatomy more accurately, and to shape how it is understood by the public at large. Importantly, the lessons learned and the model presented are directly applicable to anatomical sciences educators, offering practical pathways for integrating public scholarship into their teaching, outreach, and engagement efforts.

### Guidelines for Improving an Organization Wikipedia Article

1. **Conduct a Baseline Audit**: Evaluate the existing article against Wikipedia’s quality guidelines to identify content gaps and structural deficiencies (*assessment for learning*)
2. **Structure Content Systematically**: Organize information into clear, dedicated sections such as history, mission, governance, and outreach (*information architecture*)
3. **Source with Rigor**: Rely on high-quality, independent secondary sources to substantiate claims and ensure verifiability (*evidence-based practice*)
4. **Enhance Visual Appeal**: Incorporate elements like infoboxes and images sourced from Wikipedia Commons to succinctly present key details and improve readability (*dual coding theory*)
5. **Collaborate with Stakeholders**: Engage internal communications teams and subject experts to contribute accurate data and visual assets (*stakeholder theory*)
6. **Invite Community Feedback**: Use the article’s talk page and WikiProject banners to solicit constructive input from the Wikipedia community (*participatory communication theory*)
7. **Monitor and Update Regularly**: Keep the content current by tracking edits and updating information as new data or feedback becomes available (*continuous improvement theory*)

## ACKNOWLEDGMENTS

Thank you to those who provided feedback on the initial edits of the Wikipedia article. A special acknowledgement goes to AAA Communications Specialist Michelle Aguirre for her assistance in obtaining contemporary photographs for inclusion in the Wikipedia article. The University of Colorado Anschutz Medical Campus in Aurora is located on the traditional lands of the Cheyenne, Arapaho, Ute, and many other Indigenous nations. Forty-eight contemporary tribal nations maintain historical ties to the region now known as Colorado, and their enduring relationships with these lands are recognized and honored. This acknowledgment affirms the continued presence, knowledge systems, and contributions of Native peoples, while also recognizing the lasting impact of displacement and injustice. Respect and gratitude are extended to the Indigenous communities who continue to live, work, and thrive in this place.

## AUTHOR CONTRIBUTIONS

**Michael A. Pascoe:** Conceptualization; investigation; methodology; project administration; writing of original draft.

## BIOGRAPHY

**Michael A. Pascoe**, PhD, is an Associate Professor in the School of Medicine at the University of Colorado Anschutz Medical Campus. His primary role is the developer and deliverer of gross anatomy content within the DPT and PA curricula. His research explores the integration of technology in anatomy education. He is also interested in partnering with clinicians for donor-based anatomy education experiences.

## APPENDIX

List of inbound Wikipedia links to the American Association for Anatomy (AAA) article as of April 4, 2025 (total = 81), with additions as of April 15, 2025 shown in italics (revised total = 123).

Direct inbound links are Wikipedia pages that link straight to “American Association for Anatomy”. An indirect inbound link typically links to a previous name of the Association (e.g., “Association of American Anatomists” or “American Association of Anatomists”), which in turn sends the user to the correct page.

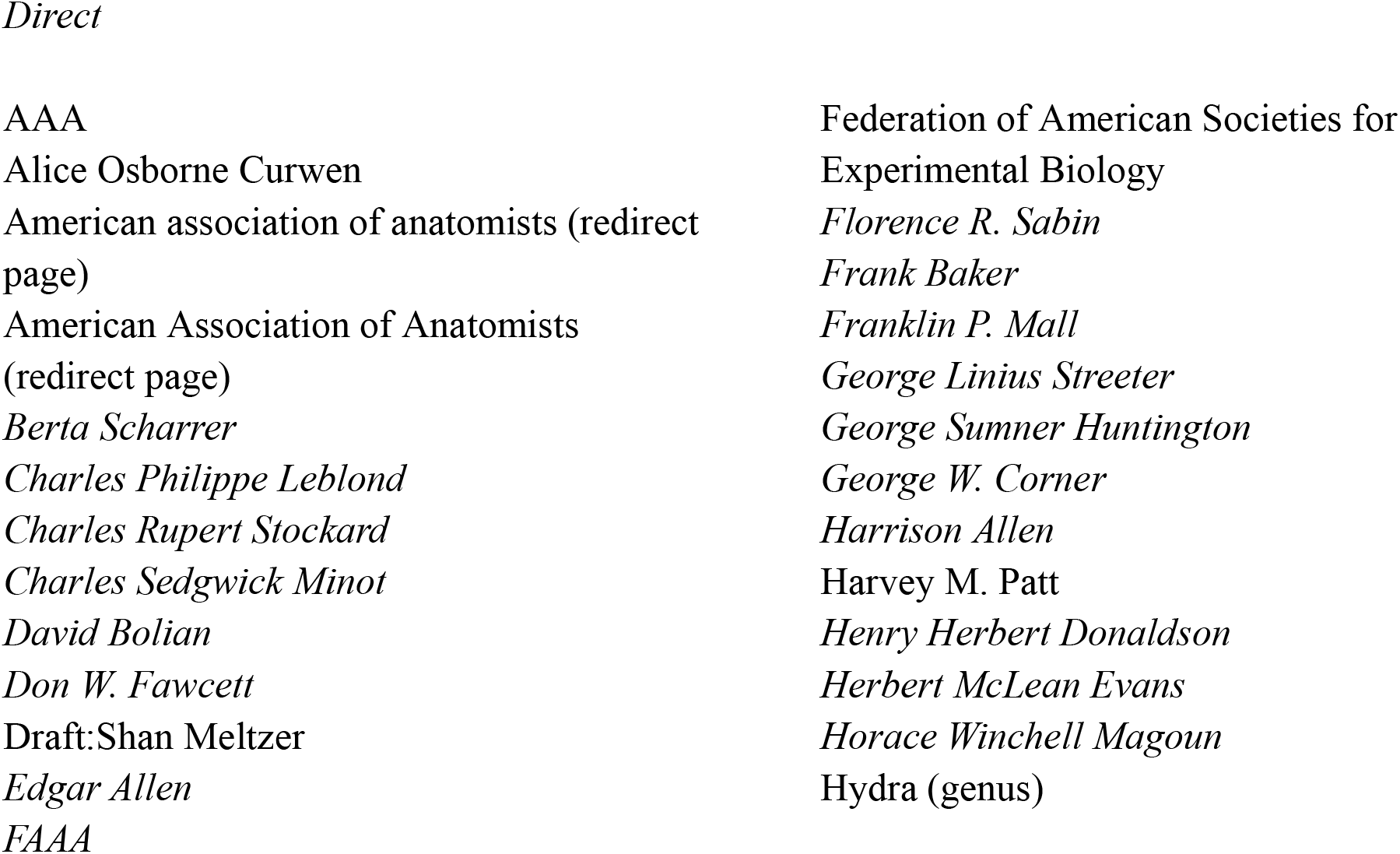
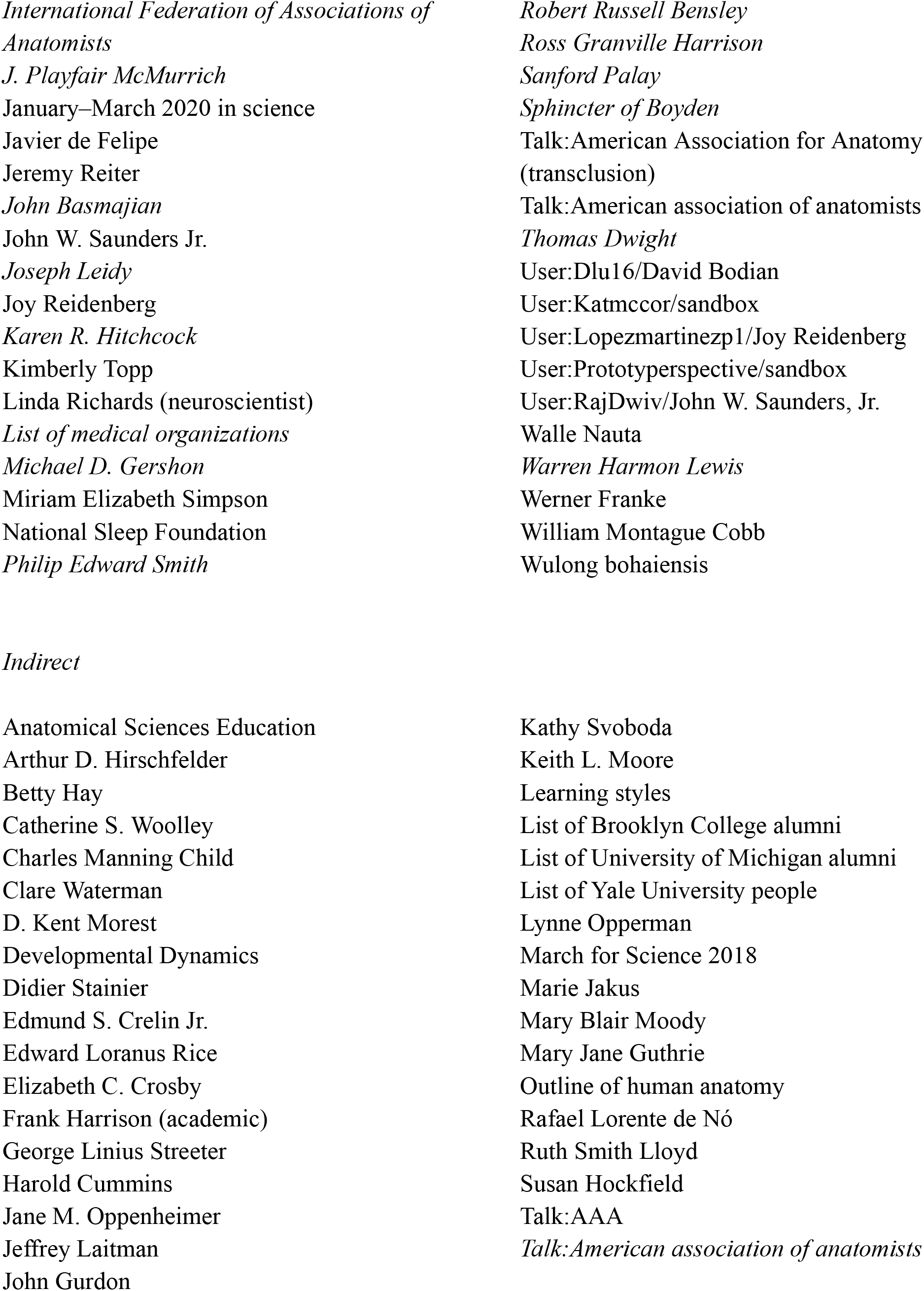
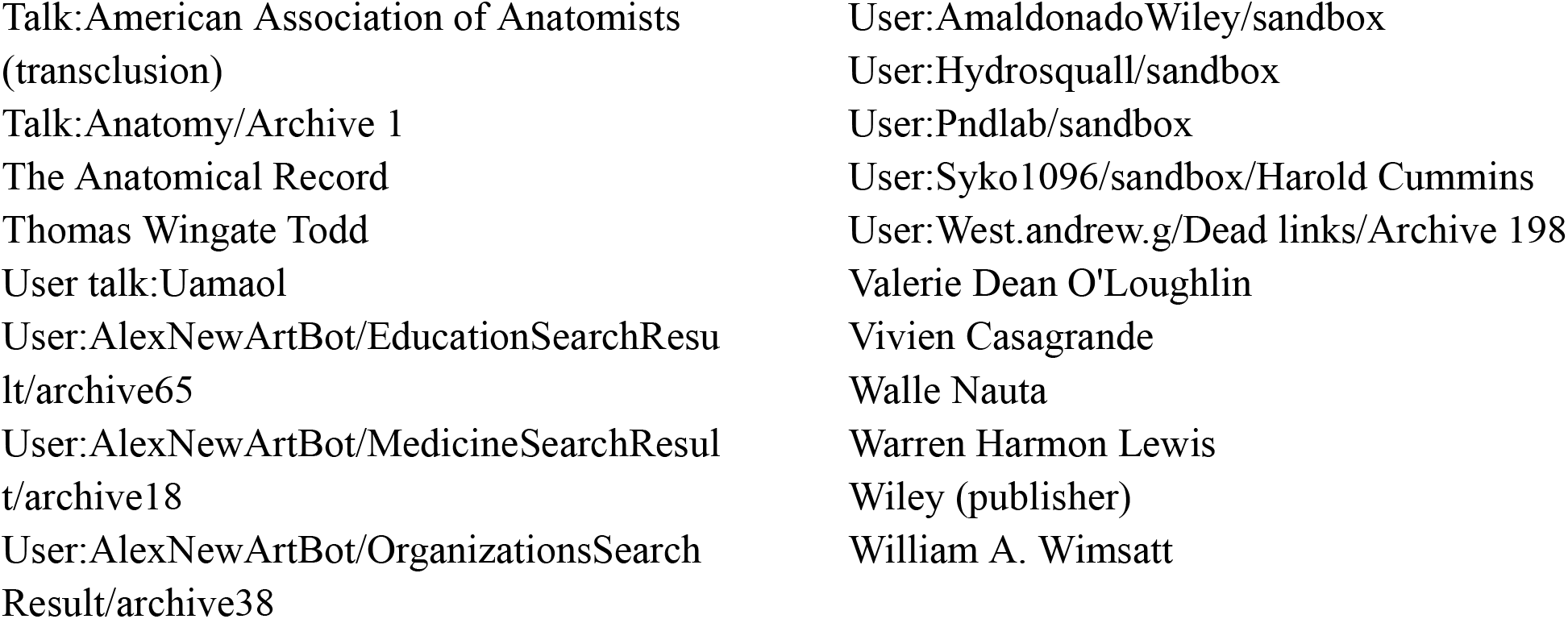

## REFERENCES

1. Peters HP. Gap between science and media revisited: Scientists as public communicators. Proc Natl Acad Sci U S A. 2013;110(Suppl 3):14102–9. doi:10.1073/pnas.1212745110.

2. Hayes DP. The growing inaccessibility of science. Nature. 1992;356(6372):739–40.

3. Bucchi M. Of deficits, deviations and dialogues: Theories of public communication of science. In: Bucchi M, Trench B, editors. Handbook of public communication of science and technology. London: Routledge; 2008. p. 71–90.

4. Pickard V, Williams AT. Salvation or folly? The promises and perils of digital paywalls. Digital Journalism. 2014;2(2):195–213.

5. National Academies of Sciences, Engineering, and Medicine. Communicating science effectively: A research agenda. Washington, DC: The National Academies Press; 2017.

6. Lih A. The Wikipedia revolution: How a bunch of nobodies created the world’s greatest encyclopedia. New York: Grand Central Publishing; 2009.

7. Laurent MR, Vickers TJ. Seeking health information online: does Wikipedia matter? J Am Med Inform Assoc. 2009;16(4):471–9.

8. Statista. Most popular websites worldwide as of November 2024, by total visits [Internet]. 2024 [cited 2025 Apr 8]. Available from: https://www.statista.com/statistics/1201880/most-visited-websites-worldwide/

9. Giles J. Special Report Internet encyclopaedias go head to head. Nature. 2005;438(15):900–1.

10. Finney D, Baleta T. For more reliable AI, academics should edit Wikipedia. Nature. 2025;639(8054):306.

11. American Association for Anatomy. Mission, vision, values [Internet]. Rockville, MD: AAA; [cited 2025 Apr 8]. Available from: https://anatomy.org/ANATOMY/ANATOMY/About-Us/mission-vision-values

12. Dunnwald M, DeLeon VB, Burrows AM. The importance of science communication and public engagement to professional associations. Anat Sci Educ. 2025. Forthcoming.

13. Xu SX, Zhang X. Impact of Wikipedia on market information environment: Evidence on management disclosure and investor reaction. MIS Q. 2013;37(4):1043–68.

14. Vincent N, Hecht B. A deeper investigation of the importance of Wikipedia links to search engine results. Proc ACM Hum-Comput Interact. 2021;5(CSCW1):1–15.

15. Wikipedia contributors. American Association for Anatomy [Internet]. Wikipedia, The Free Encyclopedia; 2024 Nov 30 [cited 2025 Apr 5]. Available from: https://en.wikipedia.org/w/index.php?title=American_Association_for_Anatomy&oldid=1260332367

16. Wikipedia contributors. Wikipedia: Manual of Style [Internet]. Wikimedia Foundation; 2023 Dec 19 [cited 2025 Apr 5]. Available from: https://en.wikipedia.org/wiki/Wikipedia:Manual_of_Style

17. Wikipedia contributors. Wikipedia: Content assessment [Internet]. Wikipedia; [cited 2025 Apr 5]. Available from: https://en.wikipedia.org/wiki/Wikipedia:Content_assessment

18. Pageview Analysis Tool [Internet]. [cited 2025 Apr 5]. Available from: https://pageviews.wmcloud.org/?project=en.wikipedia.org&platform=all-access&agent=user&redirects=0&range=all-time&pages=American_Association_for_Anatomy

19. Page Statistics Tool [Internet]. [cited 2025 Apr 5]. Available from: https://xtools.wmcloud.org/articleinfo/en.wikipedia.org/American_Association_for_Anatomy

20. Word Count Tool [Internet]. [cited 2025 Apr 5]. Available from: https://en.wikipedia.org/wiki/User:Caorongjin/wordcount

21. Wikipedia contributors. American Physiological Society [Internet]. Wikipedia, The Free Encyclopedia; 2024 Nov 30 [cited 2025 Apr 9]. Available from: https://en.wikipedia.org/w/index.php?title=American_Physiological_Society&oldid=1260330626

22. Logan DW, Sandal M, Gardner PP, Manske M, Bateman A. Ten simple rules for editing Wikipedia. PLoS Comput Biol. 2010;6(9):e1000941. doi:10.1371/journal.pcbi.1000941.

23. Apollonio DE, Broyde K, Azzam A, De Guia M, Heilman J, Brock T. Pharmacy students can improve access to quality medicines information by editing Wikipedia articles. BMC Med Educ. 2018;18:1–8.

24. Braha J. Science communication at scientific societies. Semin Cell Dev Biol. 2017;70:1–4.

